# Geographical co-occurrence of butterfly species: The importance of niche filtering by host plant species

**DOI:** 10.1101/132530

**Authors:** Ryosuke Nakadai, Koya Hashimoto, Takaya Iwasaki, Yasuhiro Sato

## Abstract

The relevance of interspecific resource competition in the context of community assembly by herbivorous insects is a well-known topic in ecology. Most previous studies focused on local species assemblies, that shared host plants. Few studies evaluated species pairs within a single taxon when investigating the effects of host plant sharing at the regional scale. Herein, we explore the effect of plant sharing on the geographical co-occurrence patterns of 232 butterflies distributed across the Japanese archipelago; we use two spatial scales (10 × 10 km and 1 × 1 km grids) to this end. We considered that we might encounter one of two predictable patterns in terms of the relationship between co-occurrence and host sharing among butterflies. On the one hand, host sharing might promote distributional exclusivity attributable to interspecific resource competition. On the other hand, sharing of host plants may promote co-occurrence attributable to filtering by resource niche. At both grid scales, we found significant negative correlations between host use similarity and distributional exclusivity. Our results support the hypothesis that the butterfly co-occurrence pattern across the Japanese archipelago is better explained by filtering via resource niche rather than interspecific resource competition.

## Introduction

Efforts to understand community assembly processes are of major importance in ecological research for obtaining past, current, and future biodiversity information. Dispersal limitations, environmental filtering via both abiotic and biotic niches, and interspecific interactions are thought to sequentially determine local community structures (reviewed by Cavender-Bares et al. 2009). Of the various relevant factors, the significance of interspecific interaction has often been assessed by examining how different species co-occur spatially even when they share similar resource niches (Diamond 1975; Gotelli and McCabe 2002). As different species with similar niches likely prefer similar environmental habitats, but may compete strongly, recent studies have often compared the significance of interspecific interactions (in terms of community assembly patterns) from the viewpoint of niche filtering, which indicates species sorting via performance and survival rates of each species against both abiotic (e.g., climate) and biotic (e.g., resource) properties from the species pool (Webb et al. 2002; Mayfield and Levine 2010).

In terrestrial ecosystems, many plant species are used by various herbivorous insects as both food resources and habitats; however, the importance of interspecific resource competition among herbivorous insects was once largely dismissed (reviewed by Kaplan and Denno 2007). Although some earlier studies described niche partitioning between co-occurring herbivores and considered that partitioning was attributable to interspecific competition (Ueckert and Hansen 1971; Benson 1978, Waloff 1979), other studies have observed frequent co-occurrence of multiple herbivorous insect species on shared host plants despite the fact that their niches extensively overlapped (Ross 1957; Rathcke 1976; Strong 1982; Bultman and Faeth 1985). Several authors have even argued that interspecific competition among herbivorous insects is too rare to structure herbivore communities (Lawton and Strong 1981; Strong et al. 1984). In subsequent decades however, many experimental ecological studies revealed that herbivorous insects do compete with one another, often mediated by plastic changes in plant defense traits following herbivory (Faeth 1986; Harrison and Karban 1986; Brown and Weis 1995; Agrawal 1999; Agrawal 2000; Viswanathan et al. 2005). These findings have prompted insect ecologists to revisit exploring the prevalence of interspecific resource competition in structuring herbivore communities (Kaplan and Denno 2007). Also, evolutionary studies have pointed out that interspecific resource competition may have regulated speciation and species diversification of herbivorous insects (Ehrlich and Raven 1964; Rabosky 2009; Nosil 2012; Thompson 2013). For example, rapid diversification after host shift to distant relative plants (Fordyce 2010) was recognized as indirect evidence of diversity regulation by interspecific competition (Ehrlich and Raven 1964; Rabosky 2009; Nakadai 2017), because the release from diversity regulation by host shift (i.e., a key innovation and ecological release; Yoder et al. 2010) may cause following specialization and diversification of herbivorous insects. In this context, the co-occurrence pattern, especially on a large spatial scale and not within the local community, is an important indicator for revealing the effects of interspecific resource competition in herbivore diversification, which eventually feeds back to their species pool as a bottom-up process (Loreau 2000). However, very little is known about the extent to which the results of the cited studies can be extrapolated to describe patterns, at the regional scale, of the distribution of herbivorous insects within a single taxon.

The effects of interspecific competition and filtering via resource niches on the co-occurrence patterns of herbivorous insects may be obscured by other potential factors. For example, climatic niches (often associated with differences in potential geographical distributions in the absence of interspecific interactions; Warren et al. 2008; Takami and Osawa 2016) may drive niche filtering, which may in turn mediate the impact of host use on assembly patterns, because the distributions of host plants are also strongly affected by climate niches (Hawkins et al. 2014; Kubota et al. 2017). In addition, taxonomic relatedness should reflect niche similarity; the niche of any organism should be partly determined by its phylogenetic history (Webb et al. 2002).

For example, phylogenetic conservatisms of host use (i.e., resource niche) in herbivorous insects were often reported (Lopez-Vaamonde et al. 2003; Nyman et al. 2010; Jousselin et al. 2013; Doorenweerd et al. 2015). Thus, the outcomes of competition within shared niches should reflect both resource and climatic factors (Mayfield and Levine 2010: Connor et al. 2013). As such factors may correlate with host use by individual species; any focus on host use alone may yield misleading results. Furthermore, dispersal ability may complicate assembly patterns, often being associated with both the extent of the geographical range and other factors (Kneitel and Chase 2004). A high dispersal ability may enhance the extent of co-occurrence. Thus, when attempting to explore the effects of competition and filtering via resource niches on distribution patterns, it is important to consider all of taxonomic relatedness, climatic niche preferences, and dispersal ability.

In the present study, we explored whether interspecific competition or niche filtering better explained the geographical co-occurrence of a group of herbivorous butterflies. Given that the host specificity of butterflies is relatively high among herbivorous insects (Novotny et al. 2010), butterflies that share a host plant are more likely to be potential competitors for each other than other less specialized herbivores are. Butterfly larvae often consume huge amounts of leaves in the larval stage, and even eat whole material of plants in some cases (Inouye and Johnson 2005; Fei et al. 2016; Hashimoto and Ohgushi 2017). Moreover, it is suggested that not all plants or parts of a plant are suitable for butterfly larvae due to within- and among-individual variations in plant quality (e.g., nutritional status and defensive traits such as secondary metabolites) (Damman 1989; Dempster 1997; Van Zandt and Agrawal 2004). Hence, the observation that most parts of terrestrial plants seem to remain unconsumed (Hairston et al. 1960; Polis 1999) does not necessary mean that butterflies are not facing resource competition. Several studies have examined butterfly–butterfly competition (Brower 1962; Millan et al. 2013), and many have reported interspecific resource competition among herbivorous insects that involved butterfly species (Agrawal 1999; Redman and Scriber 2000; Agrawal 2000; Van Zandt and Agrawal 2004; Prior and Hellman 2010). Thus, resource-mediated competition is expected to be prevalent in herbivore butterflies. In particular, we focused on herbivorous butterflies of the Japanese archipelago for the following reasons. First, the Biodiversity Center of Japan has extensive records of butterfly distribution. Second, forewing length (easily measured on photographs) is a useful index of butterfly dispersal ability (Chai and Srygley 1990; Shirôzu 2006). Finally, and most importantly, a great deal is known about the hosts used by butterfly species (Saito et al. 2016). Thus, data on Japanese butterflies can be used to explore the effects of host plant sharing and other factors on the co-occurrence patterns at regional scales.

We expected to discern one of two patterns when evaluating the significance of interspecific competition in terms of the geographical co-occurrence of herbivore butterfly. If interspecific resource competition is sufficiently intense to cause competitive exclusion with the precondition that intraspecific competition is comparably low, sharing of host plants would be positively associated with exclusive distribution (Hypothesis 1 in Table 1). Alternatively, if filtering via a resource niche is relatively stronger than resource competition, sharing of host plants would promote species co-occurrence (Hypothesis 2 in Table 1). To distinguish between these two patterns, we examined the correlations between host use similarity (i.e., the extent of sharing of host species) and the exclusiveness of geographical distribution among pairs of entirely herbivorous Japanese butterflies. The former pattern predicts that a positive correlation would be evident between host use similarity and the exclusiveness of geographic distribution. The latter pattern predicts that the correlation would be negative. Furthermore, we also considered taxonomic relatedness, climatic niche similarities, and overall dispersal abilities in the course of our work as factors affecting the association between host sharing and geographical co-occurrence (major hypotheses are summarized in Table 1).

**Table 1.**
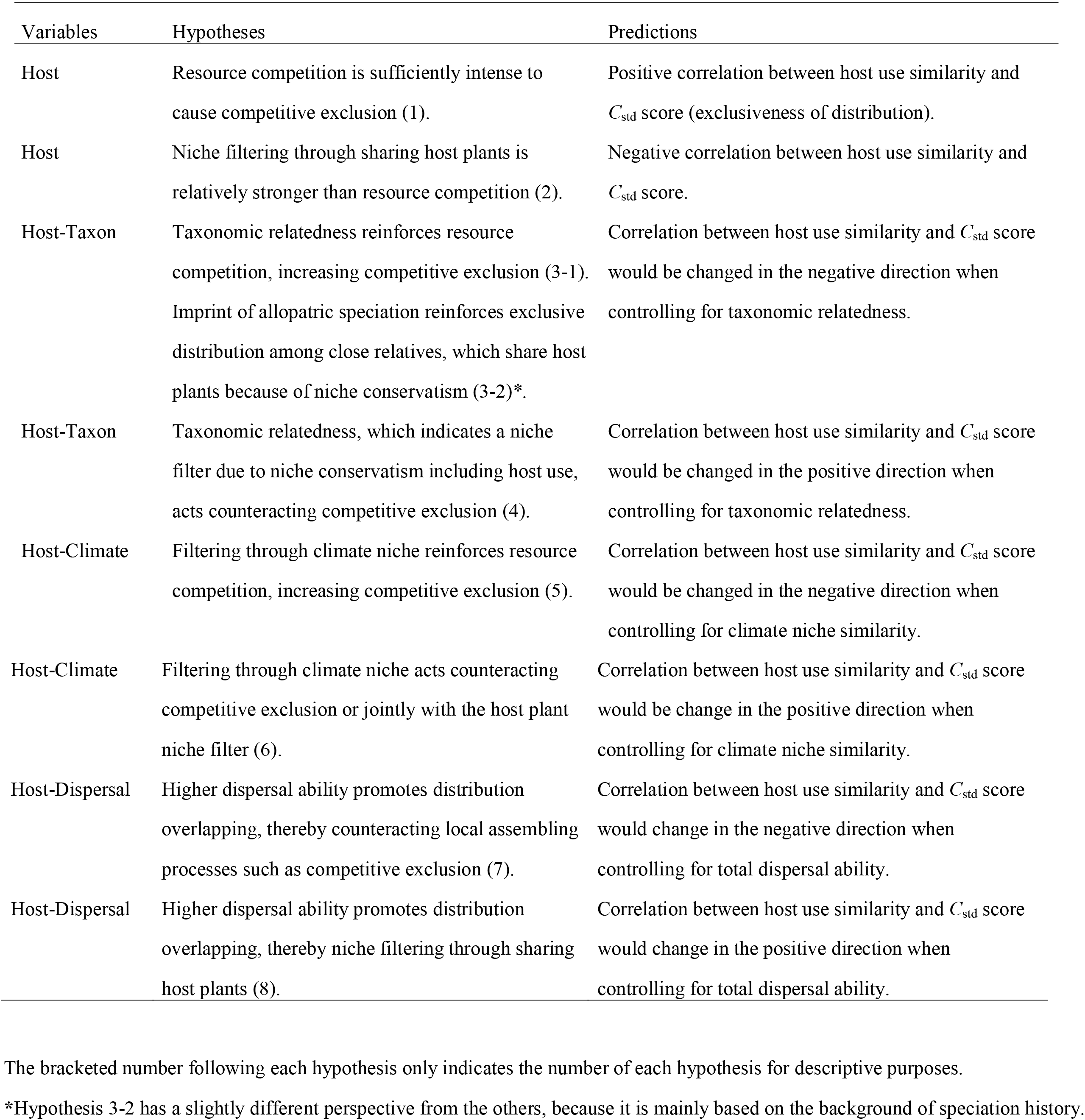
Major hypotheses and predictions tested in this study, associated with host use similarity (Host), taxonomic relatedness (Taxon), climate niche similarity (Climate), and total dispersal ability (Dispersal).

| Variables | Hypotheses | Predictions |
| --- | --- | --- |
| Host | Resource competition is sufficiently intense to cause competitive exclusion (1). | Positive correlation between host use similarity and $C_{std}$ score (exclusiveness of distribution). |
| Host | Niche filtering through sharing host plants is relatively stronger than resource competition (2). | Negative correlation between host use similarity and $C_{std}$ score. |
| Host-Taxon | Taxonomic relatedness reinforces resource competition, increasing competitive exclusion (3-1).<br>Imprint of allopatric speciation reinforces exclusive distribution among close relatives, which share host plants because of niche conservatism (3-2)*. | Correlation between host use similarity and $C_{std}$ score would be changed in the negative direction when controlling for taxonomic relatedness. |
| Host-Taxon | Taxonomic relatedness, which indicates a niche filter due to niche conservatism including host use, acts counteracting competitive exclusion (4). | Correlation between host use similarity and $C_{std}$ score would be changed in the positive direction when controlling for taxonomic relatedness. |
| Host-Climate | Filtering through climate niche reinforces resource competition, increasing competitive exclusion (5). | Correlation between host use similarity and $C_{std}$ score would be changed in the negative direction when controlling for climate niche similarity. |
| Host-Climate | Filtering through climate niche acts counteracting competitive exclusion or jointly with the host plant niche filter (6). | Correlation between host use similarity and $C_{std}$ score would be change in the positive direction when controlling for climate niche similarity. |
| Host-Dispersal | Higher dispersal ability promotes distribution overlapping, thereby counteracting local assembling processes such as competitive exclusion (7). | Correlation between host use similarity and $C_{std}$ score would change in the negative direction when controlling for total dispersal ability. |
| Host-Dispersal | Higher dispersal ability promotes distribution overlapping, thereby niche filtering through sharing host plants (8). | Correlation between host use similarity and $C_{std}$ score would change in the positive direction when controlling for total dispersal ability. |

The bracketed number following each hypothesis only indicates the number of each hypothesis for descriptive purposes.
\*Hypothesis 3-2 has a slightly different perspective from the others, because it is mainly based on the background of speciation history.

## Materials and methods

### Study area

The Japanese archipelago, including the Ryukyu Islands, forms a long chain of continental islands lying off the eastern coast of Asia. The Japanese archipelago was recognized as a hotspot of biodiversity (Gerardo and Brown 1995; Mittermeier et al. 2011) and insect diversity, with about 32,000 insect species recorded (Clausnitzer et al. 2009; Tojo et al. 2017). The latitudinal range of the archipelago (22°N to 45°N) embraces hemi-boreal, temperate, and subtropical zones. The mean temperatures in the coldest and warmest months are −19.0°C and 31.5°C, respectively; the annual precipitation ranges from 867 to 3,908 mm (Kubota et al. 2014).

### Study organisms

The Japanese archipelago hosts over 280 species of butterflies of five families (the Papilionidae, Pieridae, Lycaenidae, Nymphalidae, and Hesperiidae) (Shirôzu 2006). Over 95% of the larvae of Japanese butterflies feed on plants (Honda 2005). The host plants are diverse and include both dicots and monocots (Saito et al. 2016). Lycaenid butterflies include two non-herbivorous species, but all species of all other families are exclusively herbivorous.

### Metadata compilation

Butterfly census data are available on the website of the Biodiversity Center of Japan, Ministry of the Environment (http://www.biodic.go.jp/index_e.html). We used the results of the fourth and fifth censuses (1988 to 1991 and 1997 to 1998, respectively) of the National Survey of the Natural Environment in Japan (http://www.biodic.go.jp/kiso/15/do_kiso4.html). This database includes records of 273 species/subspecies of butterflies from the entire Japanese archipelago, in grid cells of latitude 5 min and longitude 7.5 min (the Japanese Standard Second Mesh). These grid dimensions are about 10 km × 10 km, and this grid is described below as the “10-km grid.” Furthermore, the Biodiversity Center also contains records from grid cells of latitude 30 s and longitude 45 s (the Japanese Standard Third Mesh). These grid dimensions are about 1 km × 1 km, and this grid is described below as the “1-km grid.” As processes driving community assemblies may vary between spatial scales (Cavender-Bares et al. 2009), we evaluated data from both grids. We summarized data at the species level, and converted all records into the presence or absence (1/0) of a species in each grid. Finally, the number of records used in the study after summarizing the duplicate presence records was 79,003 and 165,102 for each 10-km grid and 1-km grid for targeted species, respectively. We used the taxonomy of Shirôzu (2006).

Data on host plants and forewing length were evaluated as possible variables explaining, respectively, host use and dispersal ability. The host plants of 278 butterfly species/subspecies (3,573 records in total, after excluding duplications) were obtained from data by Saito *et al*. (2016), and the host records were described in 8 major illustrated reference books, 2 checklists, and 14 other pieces of literature; the presence of larvae on plants, and the observations of larvae eating plants or insects in the field were considered host records. We used 2,939 records of targeted species in this study (Table S1). The datasets of Saito *et al*. (2016) included some incomplete records (e.g., *Asarum* sp. of *Luehdorfia japonica, Cotoneaster* spp. of *Acytolepis puspa*) including 34 records (about 1.14% of whole records) on targeted species; those incomplete records were excluded from our analyses. Dispersal ability was evaluated by reference to adult wing length. We compiled wing data on 284 species using published illustrations (Shirôzu 2006). We used Image J software (Abramoff et al. 2004) to extract forewing lengths (in cm) from plates that included centimeter scale bars. Multiple forewing lengths were extracted when individual, sexual, and/or geographical variations were evident.

We assembled data on the distributions, host plants, climatic niches (described below), and forewing lengths of 232 butterfly species. Twenty-four species were excluded from the analysis, for the following reasons. First, the taxonomic status of three species (*Papilio dehaanii, Pieris dulcinea*, and *Eurema hecabe*) changed in the interval between the fourth and fifth biodiversity censuses (Inomata 1990; Shirôzu 2006). Thus, we excluded these species because identifications were unreliable. Second, we excluded three species of non-herbivorous butterflies (*Taraka hamada, Spindasis takanonis*, and *Niphanda fusca*). Finally, we excluded a further 18 species because the models used to evaluate their ecological niches failed to satisfy the criteria that we imposed (i.e., shortage of occurrence records for the niche modeling program or less than 0.7 of the values of area under the curve [AUC], the details are in Supplementary File 1).

### Data analysis

#### Species distribution exclusiveness

We used the checkerboard scores (C-scores; Stone and Roberts 1990) to evaluate the exclusivity of distributions between each species pair. We set *r*_i_ and *r*_j_ as the numbers of grids in which species *i* and *j*, respectively, were present; the checker unit *C*_ij_ associated with the two species was defined as: *C*_ij_ = (*r*_i_ – *S*_ij_) × (*r*_j_ – *S*_ij_), where *S*_ij_ indicates the extent of co-presence (i.e., the number of grid cells shared by the two species). Thus, the checker unit became larger as the two species occurred more commonly in different grid cells. We simulated null models to allow the observed checker units to be compared with stochastic distributions. We used the method of Jonsson et al. (2001) to maintain the observed frequencies of species occurrence and randomized the presence/absence matrices for each pair of butterfly species. The null models were run 999 times for each species pair. *C*_obs_ and *C*_null_ were the checker units of the observed and null distributions, respectively; the checker unit was standardized as: *C*_std_ = (*C*_obs_ – *C*_null_)/*SD*_null_, where *SD*_null_ indicates the standard deviation of all checker units of the null models. The checker unit of the null model, *C*_null_, was the average checker unit of all null models. Thus, positive and negative values of *C*_std_ indicate that two species are allopatrically and sympatrically distributed, respectively, to extents greater than indicated by the null models. We also attempted to construct a more constrained null model by weighting each grid by its species richness to consider the potential size of the butterfly community as well as the observed frequency of each species. However, this weighted index was very highly correlated with *C*_std_ (*r* = 0.97, *n* = 300 randomly selected pairs); thus, we adopted Jonsson’s method in this study. All statistical analyses were performed with the aid of R software version 3.2.0 (R Core Team 2015).

#### Climatic niche similarities

We used ecological niche modeling (ENM) (Franklin 2010) to evaluate climatic niche similarities among butterfly species (Warren et al. 2008). ENM associates distributional data with environmental characteristics, thus estimating the response functions of, and the contributions made by, environmental variables. Furthermore, potential distributional ranges may be obtained by projecting model data onto geographical space. In the present study, potential distributional ranges estimated by ENM should be influenced by abiotic environmental variables alone (climate and altitude); we did not consider interspecific interactions among butterflies or dispersal abilities in this context. Thus, comparisons of potential distribution patterns estimated by ENM allow evaluation of climatic niche similarities among butterfly species. The maximum entropy algorithm implemented in MaxEnt ver. 3.3.3e software (Phillips et al. 2006; Phillips and Dudík 2008) was employed in ENM analyses (Supplementary File 1 contains the details). The logistic outputs of MaxEnt analyses can be regarded as presence probabilities (Phillips and Dudík 2008). Finally, we then used Schoener’s (1968) *D* statistic to calculate climatic niche similarities between pairs of butterfly species, based on the MaxEnt outputs. *P*_x,i_ and *P*_y,i_ were the probabilities (assigned by MaxEnt) that species *x* and *y* would mesh to the extent of *i* on a geographic scale; the climatic niche similarity between the two species was defined as: *D*_env_ = 1 – 0.5 × Σ|*P_x,i_* – *P_y,i_*| where *D*_env_ ranged from 0 (no niche overlap) to 1 (completely identical niches). The probability assigned to the presence of species *x* in grid *i* was *P_x,i_* = *p_x,i_*/Σ*p_x,i_*, where *p_x,i_* was the logistic Maxent output for species *x* in grid *i*.

#### Explanatory variables

We evaluated both host use similarity and other factors that might explain exclusive species distributions (i.e., taxonomic relatedness). We calculated the total dispersal abilities of species pairs and climatic niche similarities (as explained above). Host use similarity was calculated as 1 minus Jaccard’s dissimilarity index (Koleff et al. 2003) when host plant species were shared by two butterflies. The taxonomic relatedness of each species pair was classified as: 2: in the same genus; 1: in the same family; and 0: in different families. The total dispersal ability was calculated as the sum of the ln (forewing lengths) of each species pair.

#### Statistical tests

We used the Mantel test, in which the response matrix yielded pairwise *C*_std_ data, and which included explanatory matrices, to examine the effects of host use similarity and other factors (i.e., taxonomic relatedness, climatic niche similarity, and total dispersal ability) on the exclusivity of butterfly distribution. We calculated Spearman’s correlations because one of the explanatory variables (taxonomic relatedness) was a rank variable. *P*-values were determined by running 9,999 permutations. In addition, when analyzing correlations between host use similarity and other explanatory matrices, we ran Mantel tests with 9,999 permutations, and calculated Spearman’s correlations. We used partial Mantel tests, in which the response matrix yielded pairwise *C*_std_ data, and in which a matrix of host use similarity including the other three explanatory matrices served as a co-variable, to evaluate the effects of confounding factors associated with host use similarity on the exclusivity of butterfly distribution. We ran 9,999 permutations and calculated Spearman’s correlations. We employed the vegdist function of the vegan package (Oksanen et al. 2015) and the mantel function of the ecodist package (Goslee and Urban 2007) implemented in R. We performed all analyses using data from both the 10-km and the 1-km grids. The same analysis was conducted for the datasets after excluding the species (the bottom 10% in the 10-km grid and 1-km grid scales, to reduce the bias of rare species in the analysis. The histogram of presence records for each butterfly species is shown in Figure S1.

## Results

The standardized checkerboard scores (*C*_std_ values) of most species pairs were negative, indicating that, in general, Japanese butterflies were more likely to co-occur than expected by chance (Fig. 1). The Mantel test showed that host use similarity was significantly and negatively correlated with the *C*_std_ values at both geographical scales (Fig. 1, Table 2a); all three explanatory variables exhibited significant negative correlations with the Cstd values at both scales (Table 2a). Host use similarity was significantly and positively correlated with both taxonomic relatedness and climatic niche similarity, but we found no significant correlation with total dispersal ability (Table 2b). The partial Mantel tests showed negative correlations between the *C*_std_ values and host use similarity at both geographical scales when controlling both taxonomic relatedness and dispersal ability. The differences of the correlation coefficients from the results of Mantel tests were small, specifically less than 1% (Table 2c). In contrast, we found significant positive correlations when controlling for climatic niche similarity and all three factors in the 10-km grid dataset, in which the correlation coefficients between host use similarity and *C*_std_ score largely changed in the positive direction from the results of Mantel tests, but no significant correlation in the 1-km grid dataset (Table 2c). When we excluded rare species from the analyses, most of the results were consistent with the results of original datasets, except for the correlation between the *C*_std_ values and host use similarity under the control of climatic niche similarity in the 10-km grid dataset, which was not significant (Tables S5 and 6).

**Figure 1.**
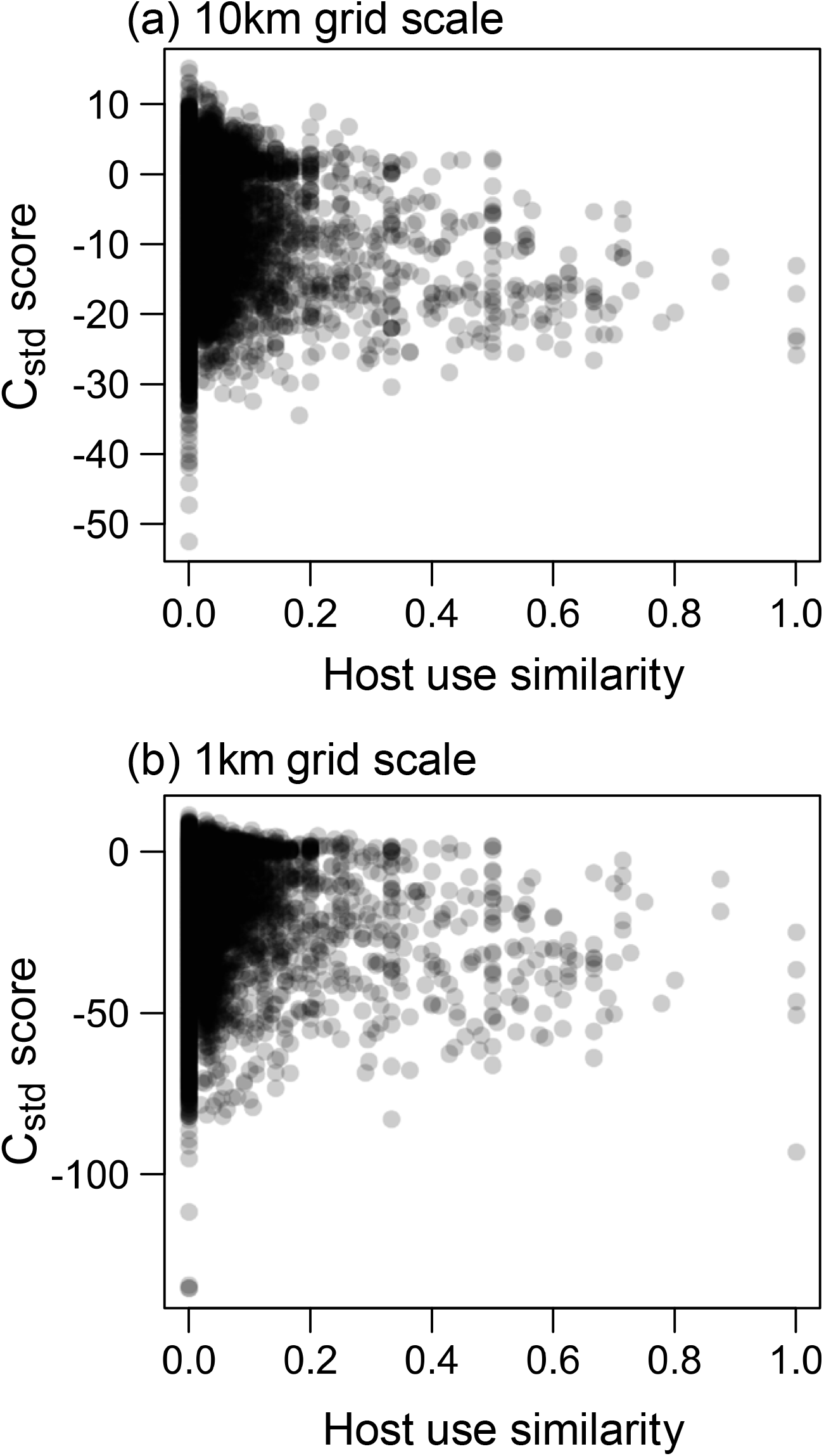
The relationships between *C*_std_ scores and host use similarities (10-km grid scale: Spearman *ρ* = −0.128, *P* = 0.0001; 1-km grid scale: Spearman *ρ* = −0.130, *P* = 0.0001; Mantel test).

**Table 2.** Correlations among host use similarity (Host), taxonomic relatedness (Taxon), climate niche similarity (Climate), and total dispersal ability (Dispersal), at two spatial scales. (a) Summary of Mantel test data on standardized *C*_std_ scores between pairs of butterfly species. (b) Summary of Mantel test data on pairwise correlations between host use similarity and the data of other explanatory matrices. (c) Summary of partial Mantel test data on standardized *C*_std_ scores between pairs of butterfly species. The “Host-Taxon” and “Host-Dispersal” data (b) were analyzed using the same datasets between two grid scales.

| (a) |  | 10-km grid |  | 1-km grid |  |
| --- | --- | --- | --- | --- | --- |
| Factor | | Mantel $\rho$ | $P$-values | Mantel $\rho$ | $P$-values |
| Host |  | -0.128 | **0.0001** | -0.130 | **0.0001** |
| Taxon |  | -0.025 | **0.0093** | -0.036 | **0.0009** |
| Climate |  | -0.718 | **0.0001** | -0.631 | **0.0001** |
| Dispersal |  | -0.045 | **0.0125** | -0.073 | **0.0008** |

| (b) |  | 10-km grid |  | 1-km grid |  |
| --- | --- | --- | --- | --- | --- |
| Factor | | Mantel $\rho$ | $P$-values | Mantel $\rho$ | $P$-values |
| Host-Taxon |  | 0.098 | **0.0001** | 0.098 | **0.0001** |
| Host-Climate |  | 0.209 | **0.0001** | 0.225 | **0.0001** |
| Host-Dispersal |  | 0.061 | <u>0.0947</u> | 0.061 | <u>0.0947</u> |

| (c) |  | 10-km grid |  | 1-km grid |  |
| --- | --- | --- | --- | --- | --- |
| Factor | Covariate | Mantel $\rho$ | $P$-values | Mantel $\rho$ | $P$-values |
| Host | Taxon | -0.126 | **0.0001** | -0.127 | **0.0001** |
| Host | Climate | 0.033 | **0.0044** | 0.016 | 0.1889 |
| Host | Dispersal | -0.125 | **0.0001** | -0.126 | **0.0001** |
| Host | All three | 0.036 | **0.0021** | 0.021 | <u>0.0890</u> |

## Discussion

Significant negative correlations were clearly evident between the *C*_std_ scores and host use similarities at both grid scales (Fig. 1, Table 2a), indicating that a pair of Japanese butterflies sharing host plants is more likely to co-occur (supporting Hypothesis 2). Significant negative correlations between *C*_std_ scores and host use similarities were evident after controlling for each of the other potentially confounding factors, taxonomic relatedness, and total dispersal ability (Table 2c). The predicted pattern from interspecific resource competition (i.e., positive correlations between the *C*_std_ score and host use similarity) were confirmed only after controlling for the effects of climatic niche similarity and all three factors in the 10-km grid data (supporting Hypothesis 6). However, the correlation coefficient was low (Table 2c), and we could not confirm the pattern in the analysis excluding rare species (Table S5c). Note that we need to interpret the results carefully, as interspecific resource competition is not the only process that generates negative correlations between *C*_std_ score and host use similarity (e.g., speciation, Warren et al. 2014). Our results are consistent with the idea that interspecific resource competition is too weak to organize communities of herbivorous insects effectively (Lawton and Strong 1981; Strong et al. 1984; Nakadai and Kawakita 2017). Rather, our results suggest that the geographic pattern of species co-occurrence among Japanese butterflies is better explained by niche filtering.

The most likely explanation of our data is that the relative strength of structuring via resource competition may be weaker than that associated with niche filtering. As the geographical distributions of host plants would be expected to be strongly associated with the local climatic environment, the impacts of resource and climatic niche filtering may combine to ensure that butterfly species sharing host plants assemble in the same places. In addition, the dispersal of adult butterflies from the patches in which they were born may counteract the structuring force imposed by interspecific competition. However, although co-occurrence was facilitated by the overall total dispersal ability (Table 2a), the negative correlations between the *C_std_* scores and host use similarity were evident even when we controlled for the effects of total dispersal ability (Table 2c). This means that dispersal alone may not explain the weak impact of resource competition on the co-occurrence patterns of Japanese butterflies. Other potential factors reducing the effects of interspecific resource competition may also be in play, although we did not address these topics in this study. For example, co-occurring butterfly species, which share their host plants, are also more likely to share their evolutionary histories. The accumulated time in which species co-occur (i.e., historical co-occurrence) may cause character displacement and reduce the effects of interspecific resource competition on the co-occurrence pattern after convergence (e.g., Germain et al. 2016). In the case of Japanese butterflies, there are differences in historical dispersal events, especially the timing and route to the Japanese archipelago, due to the repeated connections and disconnections between Eurasia continent and the Japanese archipelago (reviewed by Tojo et al. 2017). In addition, butterflies sharing host plants may experience similar historical dispersal events corresponding to the distribution changes of plant species. Such a historical process can generate faunal similarity along distance and climate (Nekola and White 1999; Kubota et al. 2017) and indirectly affect the pattern of herbivorous insects, which was also detected in our study (see the correlation between *C_std_* and climate niche similarity in Table 2a). Therefore, the effects of historical co-occurrence on the patterns of current geographical co-occurrence may also occur, although we did not discuss the details here.

The negative correlation evident between taxonomic relatedness and the *C*_std_ scores (Table 2a) suggests that niche filtering is in play among Japanese butterflies, given that taxonomic relatedness serves as a proxy of niche similarity including host use. Indeed, we found significant (positive) correlations between host use similarity and the taxonomic relatedness of Japanese butterflies (Table 2b), as has often been shown for other herbivorous insects (e.g., Nyman et al. 2010; Jousselin et al. 2013). These results reconfirm niche conservatisms associated with host use, which were also reported in previous studies (Lopez-Vaamonde et al. 2003; Nyman et al. 2010; Jousselin et al. 2013; Doorenweerd et al. 2015; Nakadai and Kawakita 2016). Moreover, when host use similarity was controlled using the partial Mantel test, taxonomic relatedness did not significantly affect co-occurrence at the 10-km grid scale (Table S4). These results suggest that, at least at the 10-km grid scale, the effects of taxonomic relatedness largely reflect host use similarity. Although we mainly discussed taxonomic relatedness as a niche similarity between species pairs throughout the manuscript, the correlation between geographical co-occurrence and taxonomic relatedness could also be interpreted as the imprint of allopatric speciation (Hypothesis 3-2, Barraclough et al. 1998; Warren et al. 2014). In that context, we can also interpret our results as the imprint of allopatric speciation not being strong enough to construct the co-occurrence pattern at the regional scale.

In the present study, we used ENM to evaluate the effects of climatic niche similarity on co-occurrence patterns. When we controlled for the effects of such niche similarity, the negative correlations between the *C*_std_ scores and host plant similarities disappeared at both spatial scales (Table 2c). This suggests that the explanatory power of climatic niche filtering is stronger than that of resource niche filtering. It is reasonable that host plants are also distributed in relation to climatic niches (Hawkins et al. 2014). Moreover, we also only found a positive correlation when controlling for the effects of climatic niche similarity in the 10-km grid data, which supports the hypothesis of interspecific resource competition (Hypothesis 6). This result suggests the possibility that the competitive exclusions of herbivorous insects do exist but usually cannot be observed due to masking by climatic niche filtering and other effects (Table 2c). It should be noted that, although ENM has been widely used to quantify climatic niches (e.g., Kozak et al. 2008; Warren et al. 2008; Takami and Osawa 2016) when we interpret this result, ENM data should be treated with caution (Peterson et al. 2011; Warren 2012; Warren et al. 2014). For example, Warren et al. (2014) noted that ENM always includes non-targeted factors that limit real distributions if those distributions correlate spatially with environmental predictors. Such confounding effects may cause overestimation of any positive correlation between climatic niche similarity and butterfly co-occurrence, and may also cause the effects of host use similarity to be underestimated when controlling for the effects of climatic niche similarity. The use of ENM to study species co-occurrence patterns requires careful consideration; more case studies evaluating the relative importance of climatic niches are required.

Although a number of studies have reported that resource competition involving butterfly species occur (e.g., Redman and Scriber 2000; Prior and Hellman 2010), few studies have indicated competitive exclusion or resource partitioning due to resource competition involving butterflies (Blakley and Dingle 1978; Millan et al. 2013). Some authors have argued that reproductive interference among butterflies sharing similar niches is a more plausible mechanism causing resource or habitat niche displacement than resource competition (Jones et al. 1998; Friberg et al. 2008, 2013: reviewed in Noriyuki et al. 2015; Shuker and Burdfield-Steel 2017). Friberg *et al*. (2013) suggested that habitat separation of two congeners of wood white butterflies *Leptidea sinapis* and *L. juvernica* is caused by reproductive interference. Because habitat displacement by reproductive interference is more likely to work at the local scale (Friberg et al. 2013), its effects may be difficult to observe at the regional or broader spatial scale. This does not contradict our results in which robust evidence of negative co-occurrence among host-sharing species was not detected.

We focused on butterfly species, which is one of the best-studied herbivore taxa in both ecology and evolution (Ehrlich and Raven 1964; Hanski and Singer 2001; Fordyce et al. 2010), as well as data on fundamental information (e.g., occurrence information and host use of each species) that are abundant in butterfly species. Our analysis relied on this advantage, enabling the use of huge datasets of butterfly species, and attempted to clarify the geographical co-occurrence on a larger scale. However, our study had several limitations. First, there was underrepresentation of rare species. The abundance and the range of distribution are different among butterfly species, and the records of rare species had some bias in our analysis using huge datasets. We additionally conducted analyses after excluding the species, which were confirmed in the lower number of grids, to eliminate the bias due to the unfounded presence records of rare species. As a result, most of the results showed the same trends as the results of the original datasets, so the bias of rare species may not be very large (Tables S5 and 6). The second issue was regarding the temporal fidelity of observations. We compiled datasets of field surveys over multiple years, but using single-year datasets, which are more appropriate for assessing the effects of interspecific resource competition on the patterns of geographical co-occurrence. If butterfly distributions largely change over the years, this approach would obscure the real patterns of resource-driven butterfly co-occurrence. Finally, we only used occurrence data with the accuracy of a large grid (10-km and 1-km squares); combining datasets of deferent resolution (grid-level and community-level data) may help to resolve co-occurrence patterns among herbivore butterflies (Cardillo and Warren 2016).

## Conclusions

The significance of interspecific resource competition in terms of the structuring of herbivorous insect communities is a source of long-standing controversy. Many researchers have sought to explain the general patterns of relationships between the co-occurrence of, and the use of different niches by, herbivorous insects. However, the data remain limited because most previous studies employed narrow taxonomic and spatial scales. In this context, our study is the first to provide a comprehensive picture of the co-occurrence patterns among a single taxonomic group over a large region. Although co-occurrence of Japanese butterflies is more likely to be driven by niche filtering than interspecific resource competition, it is essential to employ broad taxonomic and spatial scales when attempting to reveal general patterns of community assembly among herbivorous insects. Future studies should explore the relative importance of each assembly stage not only ecologically but also over evolutionary time (Rabosky 2009). Such work would answer the important question: “Why have herbivorous insects become one of the most diverse groups of the natural world?”

## Acknowledgements

We thank K. Kadowaki for his comments and advice on the original version of our manuscript, M. U. Saito for giving us the detail information of collecting datasets of host plants, the Biodiversity Center of Japan for allowing access to butterfly data at the 1-km grid scale, and the associate editor and three reviewers for their comments, which improved our manuscript. We are particularly grateful to the entomologists and naturalists who accumulated the information on butterflies used in this study. This work was supported by a grant from the Grant-in-Aid Program for JSPS Fellows (Grant No. 15J00601).

## Supporting Materials

**Table S1.** Japanese butterfly species analyzed in this study.

**Table S2.** Summary of the MaxEnt data at the 10-km grid scale.

**Table S3.** Summary of the MaxEnt data at the 1-km grid scale.

**Table S4.** Correlations among host use similarity (Host), taxonomic relatedness (Taxon), climate niche similarity (Climate), and total dispersal ability (Dispersal), at two spatial scales. (a) Summary of Mantel test data on pairwise correlations between the explanatory matrices. (b) Summary of partial Mantel test data on standardized *C*_std_ scores between pairs of butterfly species. The “Taxon-Dispersal” data (b) were obtained using the datasets at either grid mesh scale.

**Table S5.** Results using the datasets that excluded rare species. This table corresponds to Table 2.

**Table S6.** Results using the datasets that excluded rare species. This table corresponds to Table S4.

**Figure S1** Histogram of the number of observed grids for each butterfly species at both 10-km and 1-km grid scales: (a) all targeted butterfly species at the 10-km grid scale, (b) butterfly species at the 10-km grid scale, with an observed grid number less than 50, (c) all targeted butterfly species at the 1-km grid scale, and (d) butterfly species at the 1-km grid scale, with an observed grid number less than 50.

**Supplementary File 1.** Details of the methods used for ecological niche modeling.

